# High or Low Expectations: Expected intensity of action outcome is embedded in action kinetics

**DOI:** 10.1101/2024.02.20.581162

**Authors:** Batel Buaron, Daniel Reznik, Roy Mukamel

**Affiliations:** Sagol School of Neuroscience and School of Psychological Sciences, Tel-Aviv University; Max Planck Institute for Human Cognitive and Brain Sciences, Leipzig, Germany

## Abstract

Goal-directed actions are performed in order to attain certain sensory consequences in the world. However, expected attributes of these consequences can affect the kinetics of the action. In a set of three studies (n=120), we examined how expected attributes of stimulus outcome (intensity) shape the kinetics of the triggering action (applied force), even when the action and attribute are independent. We show that during action execution (button presses), the expected intensity of sensory outcome implicitly affects the applied force of the stimulus-producing action in an inverse fashion. Thus, participants applied more force when the expected intensity of the outcome was low (vs. high intensity outcome). In the absence of expectations or when actions were performed in response to the sensory event, no intensity-dependent force modulations were found. Thus, causality and expectations of stimulus intensity play an important role in shaping action kinetics. Finally, we examined the relationship between kinetics and perception and found no influence of applied force level on perceptual detection of low intensity (near-threshold) outcome stimuli, suggesting no causal link between the two. Taken together, our results demonstrate that action kinetics are implicitly embedded with high-level context such as the expectation of consequence intensity and the causal relationship with environmental cues.

## 1. Introduction

To successfully interact with the world, one must be able to predict the outcome of one’s own actions. It is commonly appreciated that the kinetics of a repeated action (e.g., the action’s execution force, velocity, trajectory etc.) can be highly variable. Classical motor learning theories mostly attributed such kinetic variability to factors such as neural noise or muscle fatigue (e.g., see Schmidt, Lee, Winstein, Wulf, & Zelaznik, 2018). However, recent evidence demonstrates that variability in action kinetics can also be accounted for by higher cognitive and contextual factors such as the goal of the action (Ansuini et al., 2015; Rosenbaum, Chapman, Weigelt, Weiss, & van der Wel, 2012). For example, the kinetics of the simple act of reaching for a glass are different if the subsequent action is to drink or to pour its content (the end-state comfort effect; Comalli et al., 2016). Furthermore, it has been shown that human observers are adept at detecting such subtle nuances – allowing them to infer underlying intentions (Cavallo, Koul, Ansuini, Capozzi, & Becchio, 2016).

Not only complex movements (such as reaching) have been shown to be affected by high-level contextual factors, but also simple actions such as pressing a button. For example, it was shown that participants tend to implicitly apply more force when pressing the same button, depending on whether or not this button is expected to trigger a sound (Cao, Kunde, & Haendel, 2020; Horvath, Biro, & Neszmelyi, 2018). In addition, press force was found to depend on the timing of the sensory feedback, such that participants apply more force on a button as delay time with auditory outcome increases (Cao, Kunde, et al., 2020; Neszmelyi & Horvath, 2018). Responding to sensory stimuli was found to affect action force. For example, supplementing a visual cue with a loud auditory tone has been shown to increase applied force relative to the response to a visual cue alone (Anzak, Tan, Pogosyan, & Brown, 2011). Given that most studies only compared the existence of an association (yes/no) of action with sensory consequence, it is unclear whether action kinetics (i.e. press force) are affected by properties of the expected stimulus (e.g. stimulus intensity). Furthermore, while previous studies focused on auditory stimuli, it is still unknown whether differences in press force are generalizable to other sensory modalities as well.

One possible explanation of such outcome-dependent differences in press force could represent the encoding of the expected sensory outcome. For example, the forward model (Wolpert & Miall, 1996) suggesting that the expected outcome of the action is represented in the motor system, such that when an action is executed an ‘efference copy’ of the expected outcome is sent from motor to sensory regions. Previous studies have shown that neural activity in motor regions can differentiate between different action outcomes (Eisenberg, Shmuelof, Vaadia, & Zohary, 2011; Krasovsky, Gilron, Yeshurun, & Mukamel, 2014). In addition, the EEG readiness potential, a neural marker of movement, was found to distinguish between button presses with and without expected auditory consequences (Reznik, Simon, & Mukamel, 2018). Such expectation-dependent differential activity in motor regions may in turn result in differences in kinetic features of the executed movement. Another possible explanation for outcome-dependent force differences could be reafferent information that modulates the kinetics of the action during its execution (Cao, Kunde, et al., 2020; Novembre et al., 2018). In the current study, we aim to distinguish between the contribution of prediction and reafferent information to press force modulation.

To address these questions and characterize the link between action kinetics and sensory events, we manipulated the relationship between button presses and sensory events in the auditory, tactile, and visual modalities. Specifically, we measured applied force when pressing a button while manipulating stimulus intensity, causal relationship (temporal order between action and sensory event), and predictability between the action and outcome intensity (deterministic / random). Our results support the notion that properties of predicted sensory outcome are embedded in the kinetics of the stimulus-producing action.

## 2. Materials and Methods

### 2.1 Participants

Across all three studies, we recruited 136 participants. Data from 16 participants were discarded due to technical problems, leaving a total of 120 participants for analysis (43 males, mean age 25.08, range 18-35 years; *Study 1:* 24 participants, 7 males. mean age: 25.13, range: 21-33 years; *Study 2:* 72 participants, 29 males. mean age: 25.33, range: 18-35 years; *Study 3:* 24 participants, 7 males. mean age: 24.26, range: 20-31 years). All participants were healthy, right-handed (determined by self-report) and had normal hearing and normal or corrected to normal vision. Participants were naïve to the purposes of the study. The study conformed to the guidelines that were approved by the ethical committee in Tel-Aviv University. All participants provided written informed consent to participate in the study and were monetarily compensated for their time.

### 2.2 Force measuring device

In order to measure the force applied by the participants during button presses, we used two force sensors (Honeywell FSA series; force range 0-20N) mounted on a response box (see figure 1A for device setup). The sensors were connected to analogue pins on Arduino^®^ mega2560. The force applied to the sensors was measured as a change in the output voltage read from an analogue pin (greater voltage corresponding to greater force) and calibrated offline to values in Newton using standard weights. The voltage from each sensor was read using MATLAB Support Package for Arduino Hardware, at a rate of 60Hz. This device was used to record press force in all 3 experiments.

**Figure 1:**
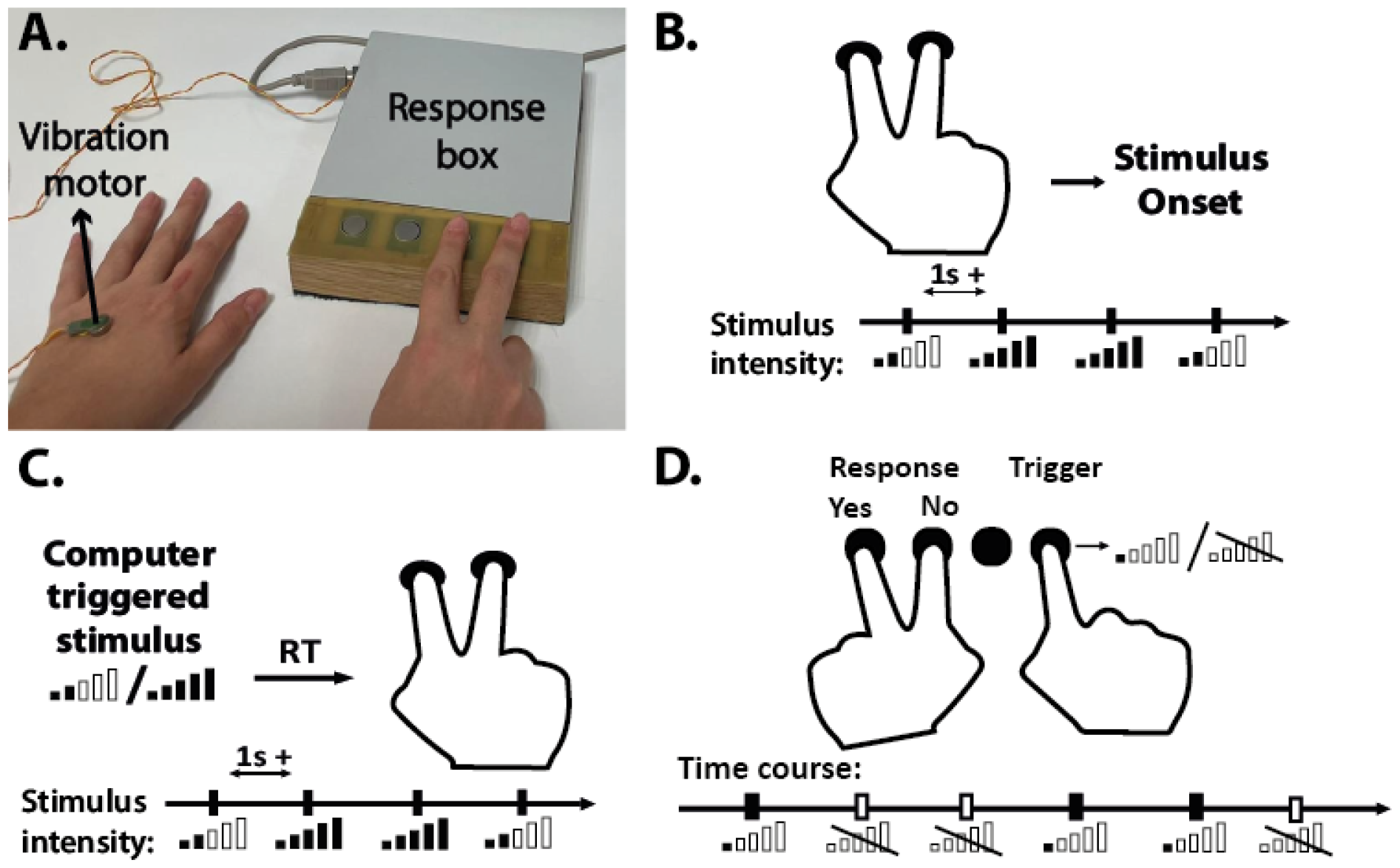
**A**. – Force measuring apparatus used in all studies and right-hand finger position (relevant to all modalities and conditions in studies 1&2) and vibration motor positioned on left hand (relevant to tactile modality in Study 2 only). **B-D:** top panels: experimental design bottom panels: experimental timeline. **B**. – Schematic illustration of experiment design for Expected and Random conditions in Study 1 and Generator condition in Study 2. Participants pressed one of two buttons (free choice) to trigger either a Low or High intensity stimulus. In the Expected (Study 1) and Generator (Study 2) conditions, participants knew in advance which button was coupled with which intensity (mapping was switched between blocks). In the Random condition (Study 1), each button triggered a tone with intensity that randomly varied across trials. **C. –** Schematic illustration of experiment design for the Follower condition (Study 2). Participants were presented with a stimulus and responded by pressing the corresponding button according to the stimulus intensity. **D. –** Schematic illustration of experiment design for the detection task (Study 3). Participants pressed a button using their right index finger that either triggered a near-threshold sound or not (random, 50% chance of triggering a sound). Sound detection was reported using one of two buttons with their left hand.

### 2.3 Hardware and software

Sounds were delivered using Creative Sound Blaster Aurora AE-5 sound card and ATH-M30x headphones (Studies 1&2) or E-A-RTONE GOLD inset air pressure earphones (Study 3). The experiment was programmed using Psychtoolbox-3 (version 3.0.16, www.psychtoolbox.org) on MATLAB 2019b (The MathWorks, Inc., Natick, Massachusetts, United States). Visual stimuli were presented using an Nvidia GTX1050TI Graphics card on a 24in screen. Vibrotactile stimulation used for Study 2 was delivered using a Shaftless Vibration Motor (size 10×3.4mm) controlled by the same Arduino board used for press force data collection.

### 2.4 Procedure

#### 2.4.1 Study 1

In order to examine the influence of expected stimulus intensity on action kinetics, participants were engaged in a sound-producing task using button-presses with either their index or middle finger of the right hand. Participants were requested to press one of two buttons to trigger a tone using either the index or middle finger of their right hand (Figure 1B). Participants were engaged in two different experimental conditions – Expected sound intensity and Random sound intensity conditions. In the Expected condition, each button was coupled either with a high or a low intensity auditory stimulus, and participants were aware of this coupling. In order to isolate force modulations that are due to prior expectations from those affected by reafferent sensory information of the evoked stimulus intensity, in a second part of the experiment, participants were requested to press the same two buttons, but on each trial the intensity of the evoked auditory stimulus was either high or low at random (Random condition). In other words, in the Random condition participants triggered the auditory stimulus by pressing the same buttons as in the Expected condition but could not predict the outcome intensity. Unbeknown to participants, we measured their applied force during button presses throughout the experiment. Importantly, in both conditions (Expected and Random), applied force had no effect on stimulus intensity (which was fixed – either high or low irrespective of applied force). The auditory stimuli were 300ms long 1000Hz pure tones, including a 15ms up and 15ms down ramping. Low and high intensity tones were fixed across participants, such that the intensity difference between the tones was 50db. At the beginning of the experiment, we verified that participants could hear the low intensity tone and that the high intensity tone was not too loud or aversive.

Throughout the experiment, participants were free to choose which button to press in each trial, however they were requested to try and balance their choices between buttons, (i.e., not to prefer one button over the other) and keep the button order as random as possible. Tone was delivered immediately when button press was detected. In order to avoid potential spill-over effects between trials, we asked participants to keep at least 1s between consecutive button presses; Trials with shorter inter-press-intervals did not trigger a tone and resulted in an error signal (color of the fixation on the screen changed to red for 300ms). Such trials were discarded from further analysis (number of errors per block: Expected condition – *M*=20.63 range: 3-64; Random condition – *M*=18.79 range: 0-64) and error trials were replaced with new ones, such that the number of valid trials was 70 for each sound intensity in each condition. The experiment consisted of 4 experimental blocks, 2 of each condition (Expected / Random). In the Expected condition, the mapping between each button and sound intensity was fixed in each block and switched between the first and second block, such that across blocks, each finger (index/middle) was mapped to high/low sound intensity. This allowed us to compare force levels across sound intensities within the same finger and avoid potential force differences between the fingers. Condition order and intensity mapping within the Expected condition were counter-balanced across participants.

#### 2.4.2 Study 2

In order to examine whether a causal relationship between actions and sensory outcome plays a significant role in the force modulations across expected stimulus intensities, we manipulated the temporal order between actions and sensory events. The experiment included two conditions - Generator and Follower – and was largely similar to the design of Study 1. The Generator condition was identical to the Expected sound intensity condition in Study 1 - participants were instructed to press buttons with known button to stimulus-intensity mappings. In the Follower condition, the temporal order (and causal relationship) between the action and stimuli was reversed. Participants were presented with either a high or low-intensity stimulus (identical to the ones used in the Generator condition) and had to respond by pressing the corresponding button (see figure 1C). Participants were requested to respond as accurately as possible with no imposed time constraint. The time interval between responses and initiation of the stimulus in the next trial was 1s. As in Study 1, each condition consisted of 2 blocks and the mapping between buttons (index/middle finger) and stimulus intensity (low/high intensity either for triggering or responding) was switched across blocks. Each block lasted until at least 70 valid trials in each condition were collected (number of errors per block in the Generator condition (too quick button presses) – M=18.674 range: 0-87; Follower condition (responding wrong intensity): mean across participants M=3.069 range: 0-20).

In order to examine whether the force modulations found in Study 1 are unique to the auditory modality, we expanded our exploration to the tactile and visual modalities as well. Participants were randomly assigned to one of the three sensory modalities, 24 participants in each modality group, such that in each modality participants completed both the Generator and the Follower conditions. In all groups, stimuli intensities were fixed for all participants. In the auditory group, auditory stimuli were identical to those used in Study 1. In the visual group, visual stimuli were Gabor patches 6° in diameter with a spatial frequency of 6 cycles per degree (cpd) located at the center of the screen. The high intensity stimulus had an 80% contrast, while the low intensity stimulus had an 8% contrast. Visual stimuli were presented for 100ms. In the tactile group, tactile stimuli were vibrations (akin to a cellular phone on vibrate mode) delivered to the back of participants’ left hand using a vibration motor controlled by an analogue pin on Arduino® mega2560 (same device used for collecting press force data; see figure 1A). High intensity vibration had a duty cycle of 0.95, while low intensity vibration had a duty cycle value of 0.42. Vibration stimulation was delivered for 300ms. Prior to the experiment, we verified that each participant could perceive the low intensity stimulus and that the high intensity stimulus was not aversive to them. Adjustments were made to stimuli if needed, but the difference between low and high intensities was kept constant for all participants.

#### 2.4.3 Study 3

In the third study we focused on the relationship between applied force and perception, examining whether changes in applied force are accompanied by changes in detection of low intensity sounds – thus alluding to a potential functional role. In this study, we examined whether detecting sounds at hearing threshold is associated with the amount of applied force used to trigger the sound. To this end, participants were engaged in a Generator task (similar to Studies 1 & 2), but this time sounds were delivered at the individual participant’s hearing threshold. Sound detection and applied force were measured.

At the beginning of the study, each participant’s hearing threshold was estimated using the ‘1 step up, 2 steps down’ method (Gelfand, 2010), with a step size of 1dB SPL, as used in our previous studies (Reznik, Guttman, Buaron, Zion-Golumbic, & Mukamel, 2021; Reznik, Henkin, Schadel, & Mukamel, 2014). Auditory stimuli were 300ms pure 1000Hz tones created using MATLAB. Each participant went through 4 rounds of threshold estimation. During each round, participants pressed a button using their right index finger to trigger a sound. Using their index and middle fingers of the left hand they reported whether or not they detected a sound. If the participant reported sound detection, on the next trial the sound intensity was lowered by 2dB. Otherwise, the sound intensity was increased by 1dB. Each round ended when the participant reported detection at a given intensity twice – and this intensity was set as the detection hearing threshold of that round. Out of the four threshold-estimation rounds, we selected the lowest sound level and verified it by presenting it to participants 10 consecutive times and examining their detection level. Sounds that were detected less than 4 times were re-examined with a sound level of +1dB, while sounds that were detected more than 7 times were re-examined using a sound level of -2dB. The converged sound intensity was used during the main experiment.

During the main experiment, participants were engaged in a Yes/No detection task in which they had to report whether they heard a sound. In each trial, participants pressed a button that triggered the auditory stimulus only in 50% of the trials. 300ms following button press, participants were presented with the question ‘Did you detect a sound?’ and had to respond as accurately as possible whether a sound was present or not using their left hand (same positioning as in the threshold detection part; see figure 1D). The experiment consisted of 6 blocks, 70 trials each (total of 420 trials across the experiment). Each block included a 50-50 ratio of randomly presented sound/no-sound trials.

### 2.5 Data analysis

In order to evaluate the applied force for triggering stimuli we computed the sum of force values (force sum) in a given time-window. We used a within-subject Student’s t test to compare the force sum between high / low intensity stimuli (Studies 1&2) and between detected and not detected sounds (Study 3). Data were analyzed using JASP (JASP Team, 2019. Version 0.16.0.0) and corrected for multiple comparisons using FDR correction (Benjamini & Hochberg, 1995). To verify enough trials for statistical comparison of force in Study 3, we excluded participants with less than 42 trials (20% of total sound trials) in either Hit or Miss trials (see results of Study 3). 9 participants (1 male) were excluded due to this criterion, leaving data from 24 participants for analysis.

## 3 Results

### 3.1 Study 1

In this study, participants pressed buttons to trigger either low or high intensity sounds. To this end, we examined the differences in force sum between low and high intensity sounds in consecutive 50ms time windows after press initiation (0-600ms after press initiation). We performed this analysis separately for the Expected and Random conditions. In the Expected condition, in which participants knew in advance which button is associated with which sound, we found a significant difference in applied force between expected low and expected high intensity sounds, such that participants applied more force (larger force sum) when they expected low intensity sound outcome relative to a high intensity sound outcome. This difference was significant for all consecutive 50ms time windows between 0-400ms after press detection / sound onset (see Figure 2A for force trajectories and Table1A for full descriptive data and statistics). In principle it is difficult to dissociate force differences that are related to prior expectation from those related to reafferent feedback of sound intensity. In order to disambiguate these two, we performed the same comparison in the Random condition, in which participants could not build prior expectation of sound intensity based on button identity. We found that in the lack of predictive knowledge, participants also applied more force when the intensity of the action outcome was low, however this effect started later than in the Expected condition and was significant between 100-300ms after press detection and sound onset (see Figure 2B for force trajectories and Table 1B for full descriptive data and statistics). No differences in press force were observed in the Random condition at the first 100ms. Significant differences after 100ms in the Random condition are probably due to sensitivity to the intensity of the auditory feedback and cannot be explained by prior expectations since evoked stimulus intensity was not known in advance. Conversely, early differences in force (less than 100ms from press onset) in the Expected condition (that are absent in the Random condition) are most likely related to the expected intensity of sound outcome. Since the aim of our study was to examine the relationship between actions and expected sensory outcome, and in order to avoid potential differences due to reafferent processing, in subsequent studies we focused our analyses only on the time window between 0-100ms. By doing so we avoided as much as possible potential contamination of effects by reafferent feedback signals.

**Table 1:**
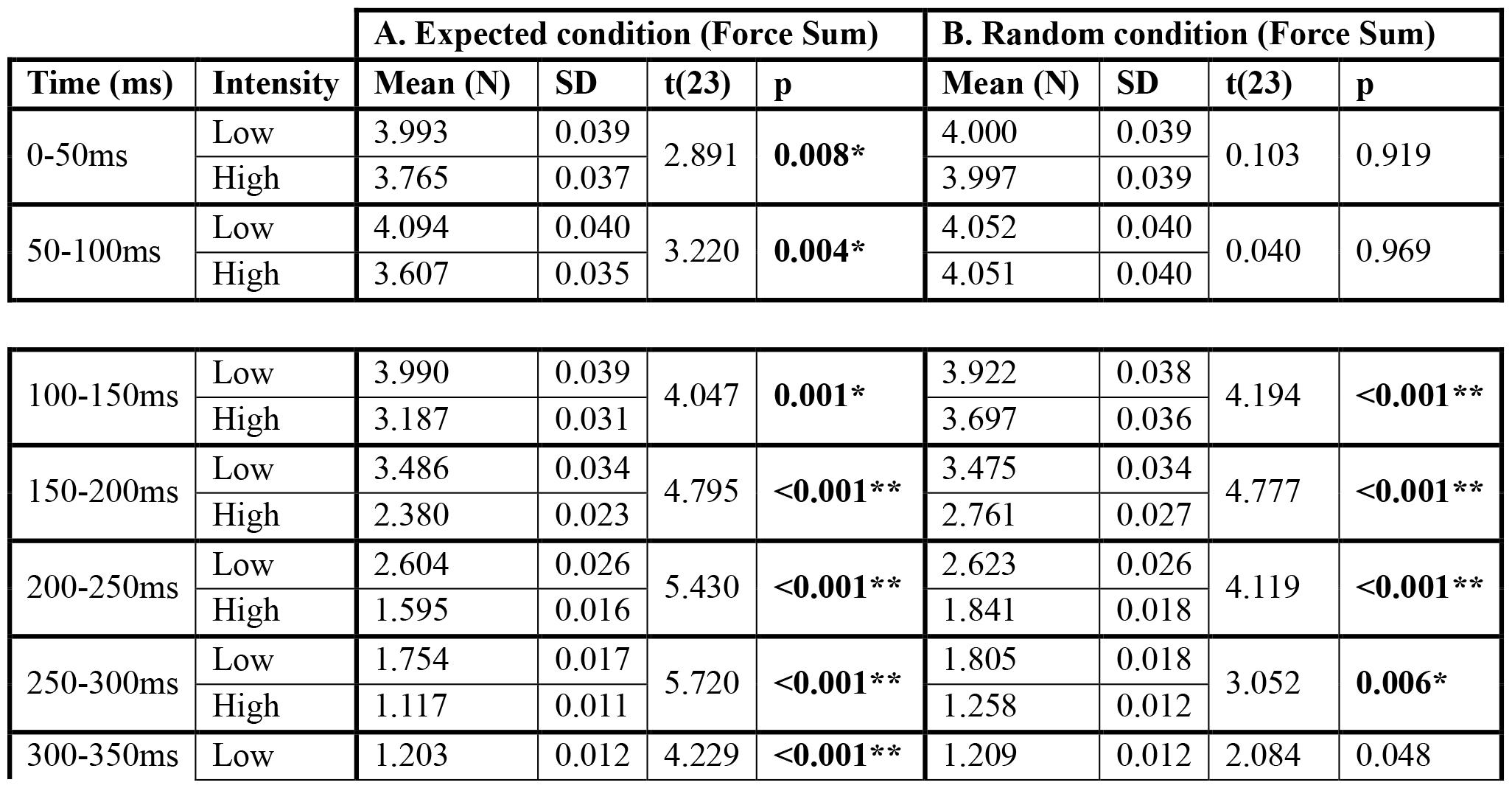
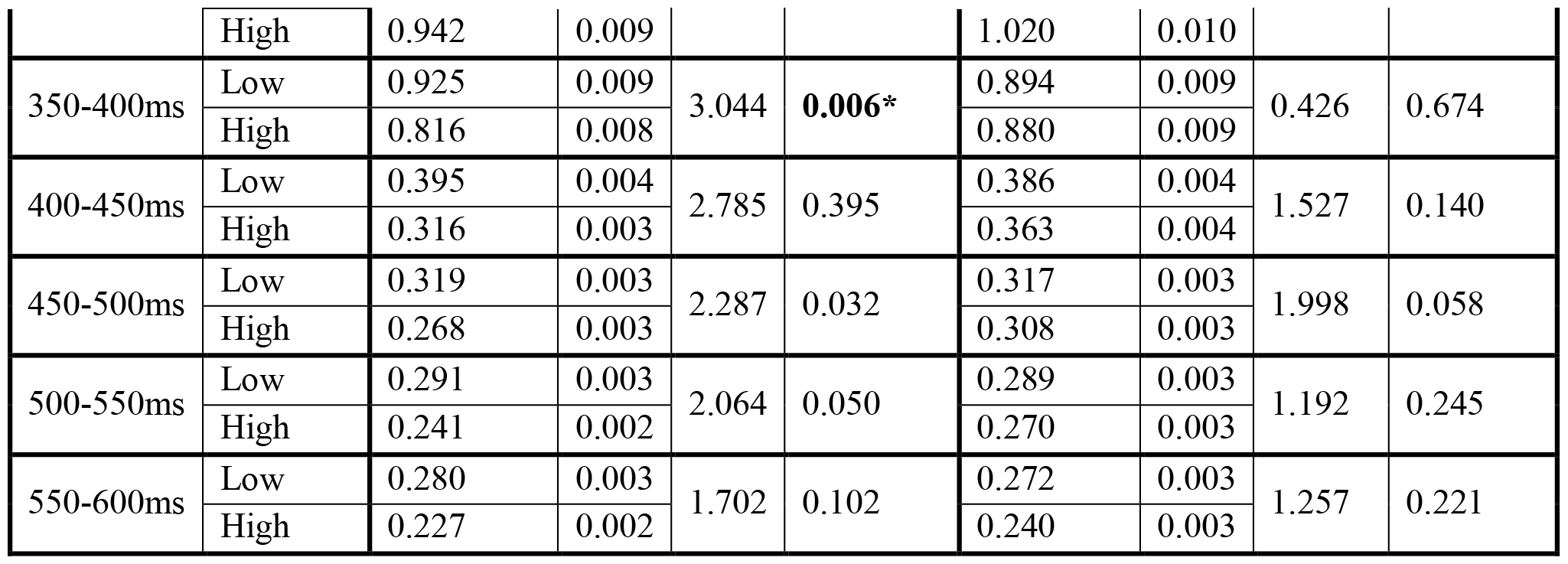
Statistical comparison of Force Sum across sound intensities and time windows in the Expected (A; left) and Random (B; right) conditions. Significant differences after correcting for multiple comparisons are marked in bold.

|  |  | A. Expected condition (Force Sum) |  |  |  | B. Random condition (Force Sum) |  |  |  |
| --- | --- | --- | --- | --- | --- | --- | --- | --- | --- |
| Time (ms) | Intensity | Mean (N) | SD | t(23) | p | Mean (N) | SD | t(23) | p |
| 0-50ms | Low | 3.993 | 0.039 | 2.891 | **0.008\*** | 4.000 | 0.039 | 0.103 | 0.919 |
|  | High | 3.765 | 0.037 |  |  | 3.997 | 0.039 |  |  |
| 50-100ms | Low | 4.094 | 0.040 | 3.220 | **0.004\*** | 4.052 | 0.040 | 0.040 | 0.969 |
|  | High | 3.607 | 0.035 |  |  | 4.051 | 0.040 |  |  |
| 100-150ms | Low | 3.990 | 0.039 | 4.047 | **0.001\*** | 3.922 | 0.038 | 4.194 | **<0.001\*\*** |
|  | High | 3.187 | 0.031 |  |  | 3.697 | 0.036 |  |  |
| 150-200ms | Low | 3.486 | 0.034 | 4.795 | **<0.001\*\*** | 3.475 | 0.034 | 4.777 | **<0.001\*\*** |
|  | High | 2.380 | 0.023 |  |  | 2.761 | 0.027 |  |  |
| 200-250ms | Low | 2.604 | 0.026 | 5.430 | **<0.001\*\*** | 2.623 | 0.026 | 4.119 | **<0.001\*\*** |
|  | High | 1.595 | 0.016 |  |  | 1.841 | 0.018 |  |  |
| 250-300ms | Low | 1.754 | 0.017 | 5.720 | **<0.001\*\*** | 1.805 | 0.018 | 3.052 | **0.006\*** |
|  | High | 1.117 | 0.011 |  |  | 1.258 | 0.012 |  |  |
| 300-350ms | Low | 1.203 | 0.012 | 4.229 | **<0.001\*\*** | 1.209 | 0.012 | 2.084 | 0.048 |
|  | High | 0.942 | 0.009 |  |  | 1.020 | 0.010 |  |  |
| 350-400ms | Low | 0.925 | 0.009 | 3.044 | **0.006\*** | 0.894 | 0.009 | 0.426 | 0.674 |
|  | High | 0.816 | 0.008 |  |  | 0.880 | 0.009 |  |  |
| 400-450ms | Low | 0.395 | 0.004 | 2.785 | 0.395 | 0.386 | 0.004 | 1.527 | 0.140 |
|  | High | 0.316 | 0.003 |  |  | 0.363 | 0.004 |  |  |
| 450-500ms | Low | 0.319 | 0.003 | 2.287 | 0.032 | 0.317 | 0.003 | 1.998 | 0.058 |
|  | High | 0.268 | 0.003 |  |  | 0.308 | 0.003 |  |  |
| 500-550ms | Low | 0.291 | 0.003 | 2.064 | 0.050 | 0.289 | 0.003 | 1.192 | 0.245 |
|  | High | 0.241 | 0.002 |  |  | 0.270 | 0.003 |  |  |
| 550-600ms | Low | 0.280 | 0.003 | 1.702 | 0.102 | 0.272 | 0.003 | 1.257 | 0.221 |
|  | High | 0.227 | 0.002 |  |  | 0.240 | 0.003 |  |  |

**Figure 2:**
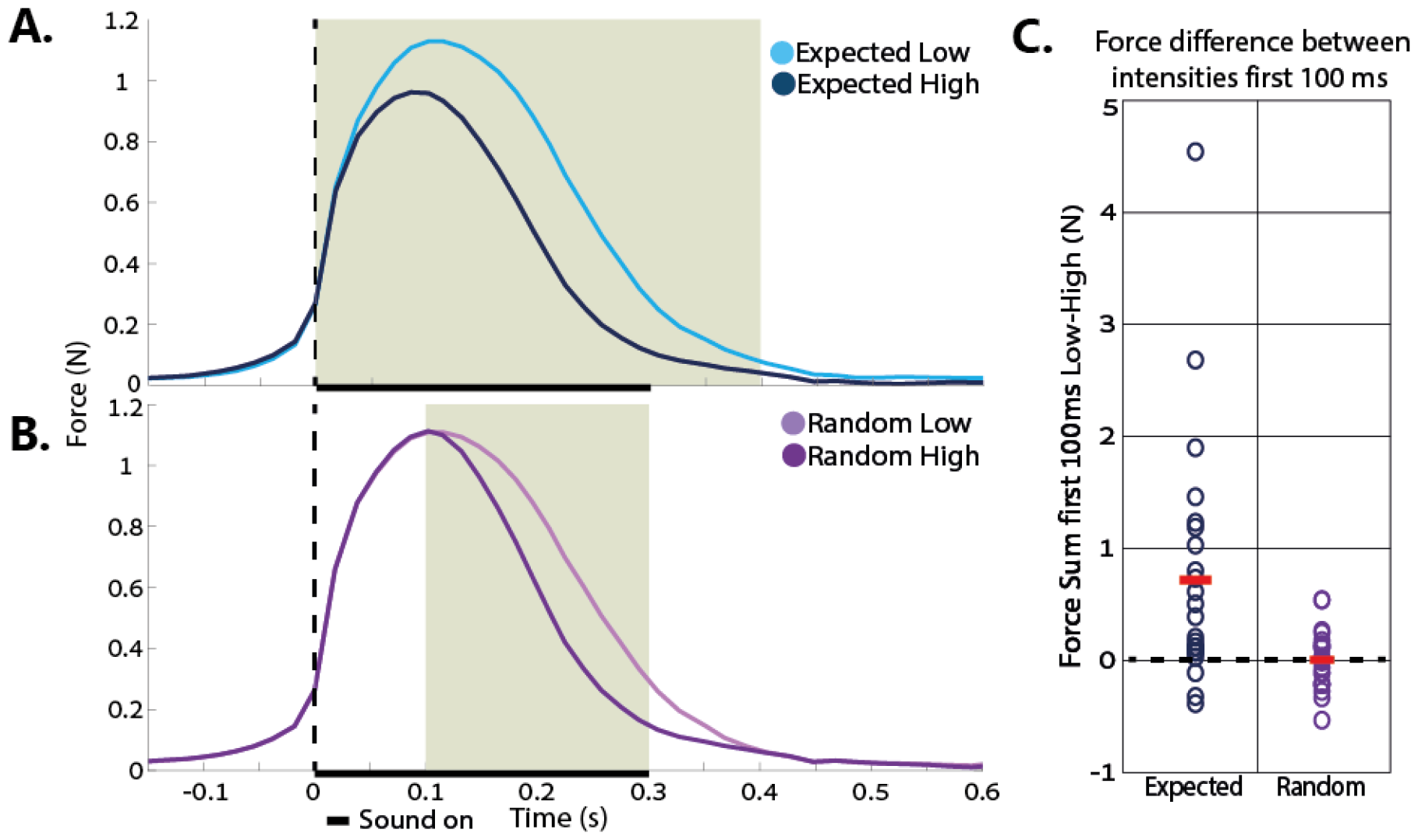
**A+B –** Force (Newton) - Time (s) trajectories for triggering Low (light colors) vs. High (dark colors) intensity sounds in the Expected (**A**) and Random (**B**) conditions. The dashed line represents press detection + sound onset (time zero). The bold line on the horizontal axis corresponds with sound duration. The shaded background represents time windows in which a significant group difference in force between stimulus intensity conditions was found (p<0.05 FDR corrected). **C –** Individual participants’ differences in force sum between Low and High intensity tones during first 100ms of button press. Red lines indicate group average, and dashed line at zero represents no difference in force sum between the two sound intensities (Expected condition: p<0.001; Random Condition: p=0.92).

### 3.2 Study 2

In order to further establish whether the differences in applied forces we found in Study 1 are related to expected intensity of action outcome, we manipulated the causal relationship (temporal order) between action and sensory events. Participants either pressed a button to trigger a sensory event (Generator condition) or pressed a button in response to a sensory event (Follower condition). Sensory events were either in the auditory, tactile or visual modalities (see methods). Based on the results from Study 1, we focused on sum force data from the time window between 0-100ms after press onset and compared the force sum in this time window between low and high intensity stimuli for each condition (Generator / Follower) and modality (Auditory / Tactile / Visual). We focused on this time window in order to examine the influence of expectations about stimulus intensity on applied force and avoid potential influence of reafferent information (see results of Study 1 for full details).

First, we performed a 2×2 repeated measures ANOVA with condition (Generator / Follower) and stimulus intensity (Low / High) as within-subjects factors, in order to compare applied force between triggering and responding to low and high intensity stimuli, irrespective of stimulus modality. We found a significant difference between conditions, such that participants applied greater force in the Follower condition (*M*=11.59 *SD*=8.61 N) relative to the Generator condition (*M*=5.45 *SD*=3.39 N; *F*(1,71)=55.29 *p*<0.001). We found a marginally significant effect of stimulus intensity, such that participants showed a tendency to apply greater force to trigger a low intensity stimulus (*M*=8.62 *SD*=7.25 N) relative to a high intensity stimulus (*M*=8.41 *SD*=7.21 N; *F*(1,71)=3.51 *p*=0.065). We also found a significant interaction effect between condition and stimulus intensity (*F*=4.44, *p*=0.039). Post-hoc test revealed that in the Generator condition there was a significant difference between low (*M*=5.65 *SD*=3.59 N) and high intensity stimuli (*M*=5.24 *SD*=3.16 N; *t*(71)=4.47 *p*<0.001), but there was no such difference in the Follower condition (Low intensity stimuli: *M*=11.60 *SD*=8.63 N; High intensity stimuli: *M*=11.58 *SD*=8.60 N; *t*(71)=0.11 *p*=0.91). See figure 3A for full force trajectories. Note that this lack of difference is not due to ceiling effect since the dynamic range of our sensor was up to 20N.

**Figure 3:**
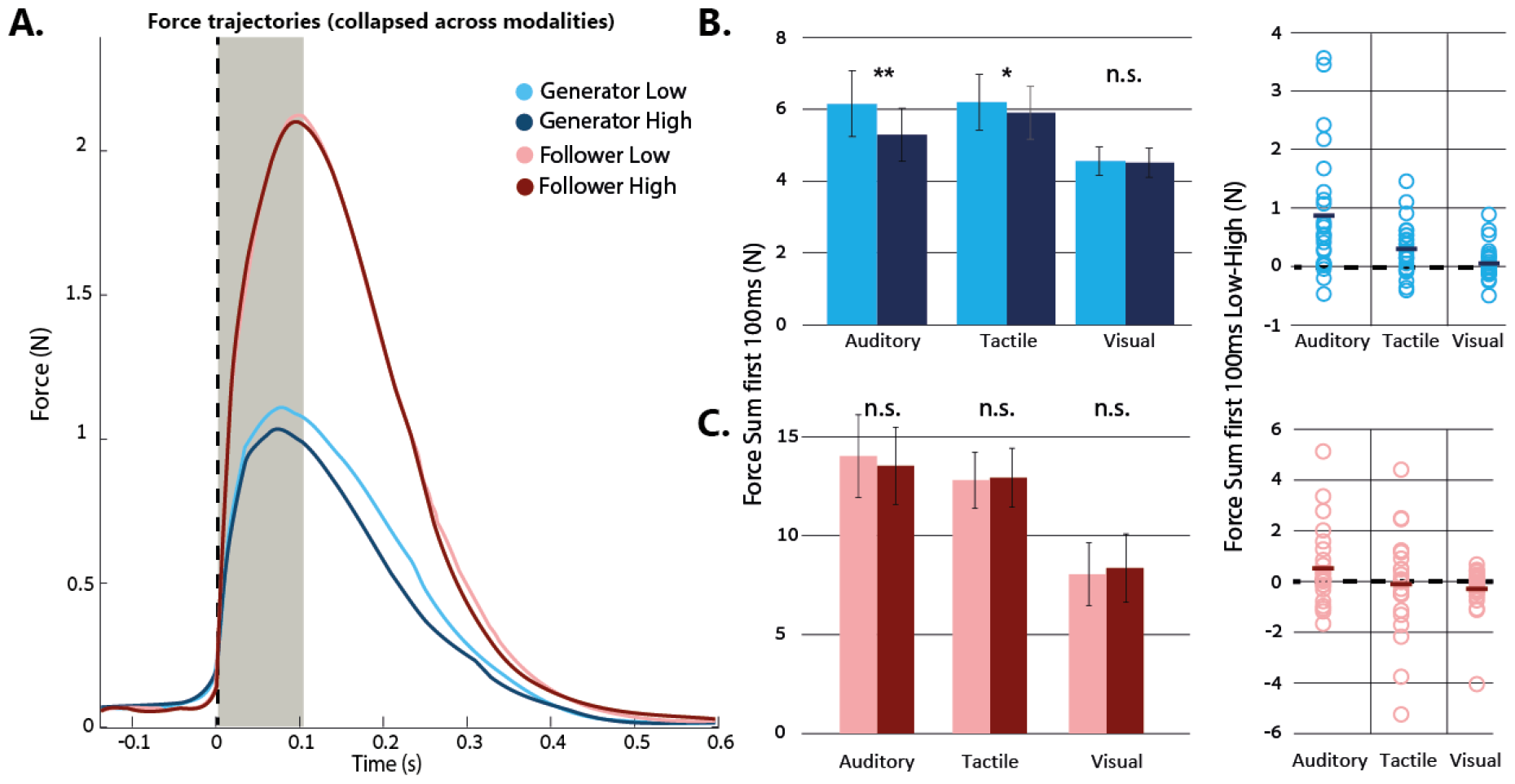
**A –**Force trajectories for triggering Low (light colors) and High (dark colors) intensity stimuli in the Generator (blue) and Follower (red) conditions. Dashed line represents press detection time (which is also stimulus onset in the Generator condition). Greyed time window marks the first 100ms used for analysis. **B –** Left panel: group mean Force Sum in the 0-100ms time window in the Generator condition, marked separately for each modality. ** p<0.001, * p<0.05. Right panel: individual participants’ differences in applied force between low and high intensity stimuli. Solid lines represent group mean difference. **C –** Same as B for the Follower condition. Note the differences in scale between the left panels in B and C (also evident in the trajectories shown in panel A).

We also examined the difference in applied force between low and high intensity stimuli within each modality separately. To this end, we used two-tailed paired sample Student’s t-test, comparing directly between Force Sum for triggering Low and High stimulus intensities within each modality and condition. Results from the Generator condition in the ***Auditory*** modality replicated the results from Study 1, demonstrating a significant difference in applied force between Low (Force Sum *M*=6.17 *SD*=4.42 N) and High intensity stimuli (Force Sum *M*=5.30 *SD*=3.55 N; *t*(23)=3.94 *p*<0.001). Similarly, in the ***Tactile*** modality, we found a significant difference in applied force between Low (*M*=6.21 *SD*=3.73 N) and High intensity stimuli (*M*=5.92 *SD*=3.55 N; *t*(23)=3.22 *p*=0.004). In the ***Visual*** modality we did not find a significant difference between stimulus intensities (Low intensity Force Sum *M*=4.57 *SD*=1.89 N; High intensity Force Sum *M*=4.52 *SD*=1.95 N; *t*(23)=0.83; *p*=0.42; see figure 3B). This pattern of results persists after applying correction for multiple comparisons. In the Follower condition, no significant differences between Low and High stimulus intensities were found across all three modalities (***Auditory***: Low intensity *M*=13.99 *SD*=10.00 N; High intensity *M*=13.50 *SD*=9.35 N; *t*(23)=1.57 *p*=0.13; ***Tactile***: Low intensity *M*=12.77 *SD*=6.80 N; High intensity *M*=12.90 *SD*=7.12 N; *t*(23)=0.31 *p*=0.76; ***Visual***: Low intensity *M*=8.03 *SD*=7.57 N; High intensity – *M*=8.34 *SD*=8.24 N; *t*(23)=1.64 *p*=0.11; see figure 3C). Taken together, these results indicate that expected stimulus intensity modulates the force participants implicitly apply when there is a causal relationship between the action and the stimulus. Such pattern of results is not observed when the causal relationship is reversed. This intensity-dependent effect on force in the generator condition is prominent in the auditory and tactile modalities but is absent in the visual modality. In the supplementary materials we present the results of an additional study in the visual domain in which we manipulated an additional visual feature (speed; Slow/Fast) of actions’ visual outcome, and also show no implicit modulation of press force (see supplementary material for full details).

Finally, we also compared the reaction time (RT) for responding to different intensity stimuli in the Follower condition. Collapsing across all modalities, participants tend to respond slower to low intensity stimuli (*M*=882.18ms *SD*=200.60ms) than to high intensity stimuli (*M*=819.59ms *SD*=204.34ms; *t*(71)=4.78 *p*<0.001). Further examining this separately in each modality, we found such effect in the Auditory (Low intensity: *M*=893.36ms *SD*=182.67ms; High intensity: *M*=795.17ms *SD*=157.26ms; *t*(23)=4.21 *p*<0.001) and Tactile (Low intensity: *M*=941.95ms *SD*=186.99ms; High intensity: *M*=883.55ms *SD*=149.76ms; *t*(23)=4.14 *p*<0.001) modalities, but no difference in RT was found in the Visual modality (Low intensity: *M*=811.24ms *SD*=208.93ms; High intensity: *M*=780.05ms *SD*=268.05ms; *t*(23)=1.14 *p*=0.26).

### 3.3 Study 3

Studies 1 & 2 point to differences in applied force that depend on the expected intensity of sensory outcome. In order to examine whether applied force affects the perception of the sensory outcome, we used an auditory detection task (see Methods). To this end, we used the force sum in the time window between 0-100ms to examine the relationship between press force and sound detection. We used median split to obtain soft / strong press trials (above / below median) and compared the signal detection theory parameters (d’ and criterion values) between strong and soft presses using a within-subjects Student’s t-test. We calculated the signal detection parameters as explained in Stanislaw and Todorov (1999). To avoid division by zero, participants with no False Alarms were assigned false alarm probability of 0.5/n where n is the number of trials. In addition, we used a Bayesian analysis to evaluate the probability of the null hypothesis for all the performed t-tests.

Compatible with our hearing threshold estimation, participants correctly reported sound detection in 52.5% of the trials in which a sound was actually generated by the button press (range across participants: 21.9 – 78.5%; Hit trials). From the trials in which button-presses did not generate a sound, the average proportion of detection reports (i.e. False Alarms) across participants was 6.1% (range: 0 – 37.6%). Force Sum values of the first 100ms of the press were split to soft and strong presses (Below median force (Soft Presses): *M*=2.749 *SD*=1.693 N; Above median force (Strong Presses): *M*=5.354 *SD*=3.016 N). For each force level separately, we calculated the sensitivity in detecting a near-threshold sound (d’) and the tendency to report sound detection (criterion). Comparing the d’ measures across force levels did not yield significant differences between presses below median Force Sum level (*M*=1.932 *SD*=0.141) and presses above median Force Sum (*M*=1.958 *SD*=0.154 z; *t*(23)=0.290 *p*=0.774; BF_01_=4.48). Similar pattern of results was found for the criterion, with no differences between presses below median Force Sum level (*M*=0.891 *SD*=0.089 z) and presses above median Force Sum (*M*=0.915 *SD*=0.089 z; *t*(23)=0.411 *p*=0.685; BF_01_=4.31; see Figure 4A).

**Figure 4:**
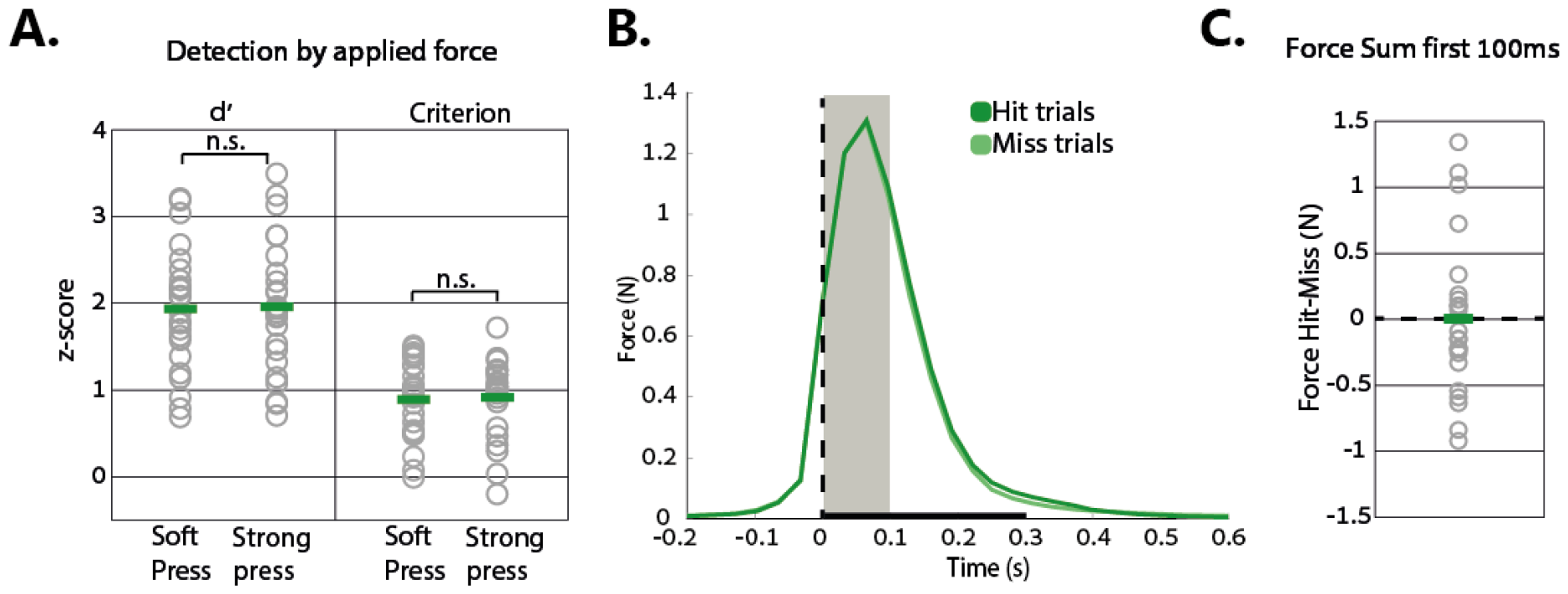
**A**. – d’ and criterion values for presses with applied force levels below and above median during the first 100ms of press. Green lines represent group mean, dots represents individual participants’ data. **B**. – Force trajectories for trials in which button-presses generated a sound separated by Hit or Miss responses. Dashed line represents press detection + sound onest. Grayed time window marks the first 100ms used for analysis. Bold line represents sound duration. **C**. – Individual participants’ differences in press force between Hit and Miss trials. Green line represents group mean difference, and dashed line at zero represents equal force applied between conditions.

Next, we directly compared the Force Sum in the first 100ms of button press between Hit and Miss trials (i.e. all trials in which the button-press generated a sound). No significant difference in applied force was found between Hit (*M*=4.020 N *SD*=2.491 N) and Miss trials (*M*=4.017 N *SD*=2.255 N; *t*(23)=0.025 *p*=0.980; see figure 4B for full force trajectory and figure 4C for mean and individual participants’ data). Further examining this null result using Bayesian analysis, we found a Bayes Factor of BF_01_=4.66, supporting the notion of no difference in press force between Hit and Miss conditions. Taken together, all our analyses point to no significant relationship between applied force level and sound detection.

## 4 Discussion

In the current study, we examined how action kinetics are affected by the properties of their coupled sensory events. To this end, we measured the implicit force levels participants apply during button presses while manipulating the buttons’ relation with sensory stimuli. We found that participants implicitly applied higher force levels when pressing a button in order to trigger a low (vs. high) intensity stimulus. We further manipulated the predictability of the outcome and the causal relationship between the action and the stimulus to evaluate their influence on press force. We found that prior expectation of stimulus intensity affects press force immediately from the onset of the action, while presses with no prior expectation started to show a difference in press force only 100ms after action onset. Furthermore, when actions followed the sensory event (i.e. no causal relationship between action and stimulus), intensity-dependent differences in force levels were abolished. Finally, we found that detection of low intensity outcome stimuli was not influenced by applied force levels, suggesting no significant functional role of force in detection of action outcome.

### 4.1 Expected intensity of action-outcome affects applied force levels

In both studies 1&2, we found an inverse relationship between the expected intensity of action consequences and applied force, such that the expectation of low intensity outcome corresponds with higher force levels. Furthermore, when button presses were not associated with expectation of an auditory outcome (as in the ‘Follower’ condition in experiment 2 in which button presses did not produce an auditory outcome), we found that participants applied the highest amount of force. Interestingly, previous studies that manipulated the existence (yes/no) of auditory outcome, rather than expectation of outcome property (intensity) report a compatible phenomenon. Participants apply less force when a button is associated with an auditory consequence and more force when the same button press was silent (Cao, Kunde, et al., 2020). This phenomenon is not specific to button presses but generalizes to other actions, such as pinches and taps (Horvath et al., 2018; Neszmelyi & Horvath, 2017). Taken together, these results point to a graded relationship between the expected intensity of auditory outcome (no outcome/low-intensity/high-intensity) and implicit force applied. The lower the expected intensity of sensory outcome, the higher the applied force.

Previous studies examining the relationship between press force and outcome focused mainly on the auditory domain (Cao, Kunde, et al., 2020; Horvath et al., 2018; Neszmelyi & Horvath, 2017, 2018), without examining whether this association generalizes to other modalities, potentially pointing to a fundamental motor mechanism. In our second study, we further examined such relation in the tactile and visual modalities. We found force differences between generating different stimulus intensities only in the auditory and tactile modalities, while it was absent in the visual modality. Lack of differences in the visual modality also persisted when examining different aspects of the visual stimuli (speed; see supplement materials). While it is plausible that in the visual modality, applied force levels encode a different parameter from contrast/speed that was not examined in the current study, our results support a functional difference in action-outcome integration in visual vs. the tactile and auditory modalities. Indications for the uniqueness of such integration in the visual modality can also be found in other paradigms. For example, in the tactile and auditory modality, it is established that self-triggered sensory stimuli are perceived as less intense relative to identical stimuli generated externally (sensory attenuation; Blakemore, Frith, & Wolpert, 1999; Kilteni & Ehrsson, 2017; Weiss, Herwig, & Schutz-Bosbach, 2011; Weiss & Schutz-Bosbach, 2012). However, in the visual modality there is relatively little consensus about the directionality of such effects (Buaron, Reznik, Gilron, & Mukamel, 2020), with some reporting an attenuation of self-triggered visual stimuli (Cardoso-Leite, Mamassian, Schutz-Bosbach, & Waszak, 2010; Dewey & Carr, 2013) while others report an enhancement of such stimuli (Desantis, Roussel, & Waszak, 2014; van Kemenade, Arikan, Kircher, & Straube, 2016). This might be a reflection of a closer association between motor and auditory/tactile modalities relative to the visual modality. Further study is needed in order to understand such differences in action-outcome integration across modalities.

### 4.2 Level of applied force depends on foreknowledge about the intensity of sensory outcome

When requesting participants to press a button to trigger a sensory stimulus, press-duration is sufficiently long (∼400ms) to be affected by both expectation processes and reafferent processing of perceived stimulus intensity. Previous studies have shown that reafferent information can affect the amount of applied force (Cao, Kunde, et al., 2020; Novembre et al., 2018). Compatible with these results, in Study 1 we show that in a Random condition, in which participants could not build an expectation of the action outcome intensity in advance, we found intensity-dependent modulations only 100ms after press initiation – presumably due to reafferent feedback. Therefore, in our analysis we focused on the first 100ms in which any force differences across conditions are more likely to be associated with motor planning and not contaminated by feedback. This time frame for reafferent information is compatible with previous results, suggesting it takes reafferent information ∼70ms to affect action force (Cao, Kunde, et al., 2020). Interestingly, the magnitude of applied force during the first 100ms in the Random condition was similar to the force used for triggering low intensity sounds in the Expected condition. This suggests that in case of uncertainty, participants behave as if they expect low intensity sound.

To further examine whether the force modulations are due to *expectation* of sensory outcome intensity, rather than an association between stimulus intensity and actions, in our second study we manipulated the temporal order between the action and the sensory event. Since expectations are associated with future events, intensity-dependent force-modulations should not be found when the stimulus precedes the action. We found that the initial level of applied force is not sensitive to stimulus intensity when the stimulus precedes the action, but only when the action is used to generate the stimulus. Note that the lack of difference in the Follower condition is not likely to be explained by a ceiling effect of press force. We measured an average of ∼2.5N applied in the Follower condition, while our force sensitive sensors are capable of measuring up to 20N. Moreover, previous studies examining press force showed that press force can be over 3N in some conditions (Cao, Kunde, et al., 2020; Neszmelyi & Horvath, 2018). The lack of differences between force intensities in the Follower condition is not in agreement with a previous study that shows an increase in press force when responding to different intensity stimuli (Ulrich, Rinkenauer, & Miller, 1998). In this study, Ulrich et al. (1998) asked participants to make a speeded reaction towards a tone delivered in 3 different intensities, showing increased force for responding to stronger sounds. Unlike in the current study, the response to the sounds was speeded, such that the task was to respond as fast as possible regardless of sound intensity. On the other hand, in the current study, the task was to cognitively report the intensity of the stimulus, such that accuracy was more important than speed. Our results indicate that in such a task, participants apply similar force levels across stimulus intensities.

### 4.3 Functional role of force modulation

Our results demonstrate an inverse relationship between expected stimulus intensity and applied force levels – with participants applying more force when the expected outcome intensity was low. Although our low intensity stimuli were well above perceptual thresholds, and participants could detect them in all of the trials, this finding raises a possible functional connection between force and perception – suggesting that participants might be applying higher force levels to facilitate perception. Previous studies have shown that perception of low intensity sounds is enhanced when those sounds are self -triggered (Reznik et al., 2021; Reznik, Henkin, Levy, & Mukamel, 2015; Reznik et al., 2014). Therefore, in our third study, we used a sound-detection task to examine the relationship between applied force and detection but found no correspondence between the two measures. This is in agreement with another study that examined whether applied force levels affect auditory discrimination, using a comparison (rather than detection) paradigm (Endo et al., 2021). In this study, participants had to press buttons at three different pre-determined force levels and their auditory discrimination was measured. They found that discrimination performance was invariant to applied force levels. Taken together, at least with respect to perception, applied force does not seem to play a significant functional role.

One possible functional role of applied force, presented by Neszmelyi and Horvath (2018), suggest it is related to the degree of association between action and outcome in an inverse fashion. In this study, they used a similar task to our ‘Generator’ condition but introduced temporal delays between the action and auditory outcome. They report increasing force levels with increasing temporal delays, reaching plateau ∼200ms at which delay the applied force levels were similar to those applied when the button press did not produce a sound (silent condition) (Neszmelyi & Horvath, 2018). Although agency was not explicitly probed, it is possible that increasing the temporal delay between action and consequences diminishes the feeling of agency over the sound and after 200ms such binding is lost. The link between agency and press force is further supported by the intentional binding task (Haggard, 2017), showing that stronger presses are associated with weaker measures of agency (Cao, Steinborn, Kunde, & Haendel, 2020). Taken together, these results suggest that press force might be associated with the binding between action and consequence and therefore represent levels of agency.

Another possible explanation for outcome-intensity dependent force modulations was presented by Kunde, Koch, and Hoffmann (2004). They suggest that when performing an action, we aim to have an “average” amount of sensory feedback, therefore actions with lower intensity outcome will be compensated by increased press force, resulting in increased tactile feedback from the press (thus compensating for the lower intensity of expected sensory feedback). Whether or not force modulations have a functional benefit or are simply an epiphenomenon related to neural processes remains to be determined.

### 4.4 Conclusion

Action kinetics are a rich measure which is influenced by various high-level cognitive constructs such as intentions and future goals. Our results show that properties of the expected sensory outcome are also embedded in applied force measures that can be observed even in a simple button-press task. Thus, by measuring subtle differences in kinetics one can infer the degree of motor-sensory binding. Such a phenomenon can be utilized in future studies as a marker for expectation and also provide a behavioral window into the neural circuits guiding behavior.

## Acknowledgements

The study was supported by the Israel Science Foundation (grant No. 2392/19 to R.M.). The authors thank lab members for constructive comments and fruitful suggestions.

