## Supplemental Data 1 for "High or Low Expectations: Expected intensity of action outcome is embedded in action kinetics"

**Supplementary materials:**

In our second study, we found a difference in press force between low and high intensity stimuli outcome in the auditory and tactile modalities. This pattern of results was not found in the visual modality, with participants applying similar levels of force for triggering low and high contrast visual stimuli. One possible explanation for such a result may be driven by the physical properties of our stimuli, such that the perceptual difference between the low and high contrast stimuli were not big enough. Another possible explanation could be that the perceptual feature we examined (contrast) was not encoded in press force, yet other physical properties may be encoded. For example, in the tactile and auditory modality there is a natural link between action force and outcome, with more vigor actions leading to stronger outcomes. In the visual modality on the other hand, there is no direct association between action vigor and visual contrast or intensity, but there is a stronger link with between vigor actions causing faster movement of visual flow. Therefore, in this study we examined these two options by creating two experimental generator conditions: Contrast and Speed. For the contrast feature, we introduced larger differences between low and high contrast stimuli from those used in study 2 described in the main text, and for the visual flow feature we introduced stimulus rotation that was either slow or fast.

**Methods:**

We recruited a new group of 24 participants (3 males, mean age = 23.25y range: 19-31y). Inclusion criteria were similar to the studies described in the main text. All participants completed two experimental conditions: triggering visual stimuli with different contrast (Contrast condition) and triggering visual stimuli rotating at different speeds (Speed condition). The Contrast condition was identical to the generator visual condition from study 2 in all parameters except we used a bigger difference between the low and high contrast (low contrast=5%, high contrast=95%) and display time of the stimuli after press was increased to 600ms. In the speed condition participants were presented with a Gabor textured circle (95% contrast) in the center of the screen and on each trial, they were requested to press one of two buttons to make the circle rotate for 600ms. Pressing one button made the circle rotate slowly (133.3°/s) while pressing the other button made the circle rotate fast (400°/s). Participants were informed in advance which button triggered which speed and were requested to freely choose which button to press and observe the stimulus rotating. As in studies 1&2, mapping between stimulus speed/contrast and buttons was reversed at the middle of the experiment and participants were informed about this reversal. The setup and force measurement device for this study was identical to the ones used in studies 1-3.

**Data analysis:**

In order to evaluate the difference in applied force for triggering visual stimuli with different contrast and different speed we used a 2-way ANOVA with condition (contrast / speed) and intensity (low contrast or speed / high contrast or speed) as within participant factors. We further directly compared between intensities within each condition using a paired sample Student's t-test and used a Bayesian analysis to further examine these results. In light of the results from study 1, all analyses were conducted on the sum of applied force in the first 100ms of action. Analysis was conducted using JASP (JASP Team, 2019. Version 0.16.0.0).

**Results:**

Examining the two-way ANOVA, we found a significant main effect of condition($F(1,23)=19.46$, $p<0.001$), with the contrast condition issuing stronger presses ($M$=4.81N $SD$=1.94N) than the speed condition ($M$=3.30 $SD$=0.83), regardless of stimulus intensity. We did not find a significant effect of intensity (Low contrast / speed: $M$=4.06N $SD$=1.22N; Hight contrast / speed: $M$=4.04N $SD$=1.26N; $F(1,23)=0.22$, $p=0.64$) or an interaction between condition and intensity ($F(1,23)=0.20$, $p=0.66$; see figure S1A). Next, we directly compared between intensities in each condition using paired sample t-test and found no differences between intensities in the contrast (Low contrast: $M$=4.81N $SD$=1.90N; High contrast: $M$=4.81N $SD$=1.99N; $t(23)=0.04$ $p=0.96$ BF(01)=4.65; replicating our main result in study 2 with higher contrast differences) or speed (Slow: $M$=3.32N $SD$=0.96N; Fast: $M$=3.27N $SD$=0.77N; $t(23)=0.52$ $p=0.60$ BF(01)=4.11) conditions (see figure S1B for individual subjects differences) .

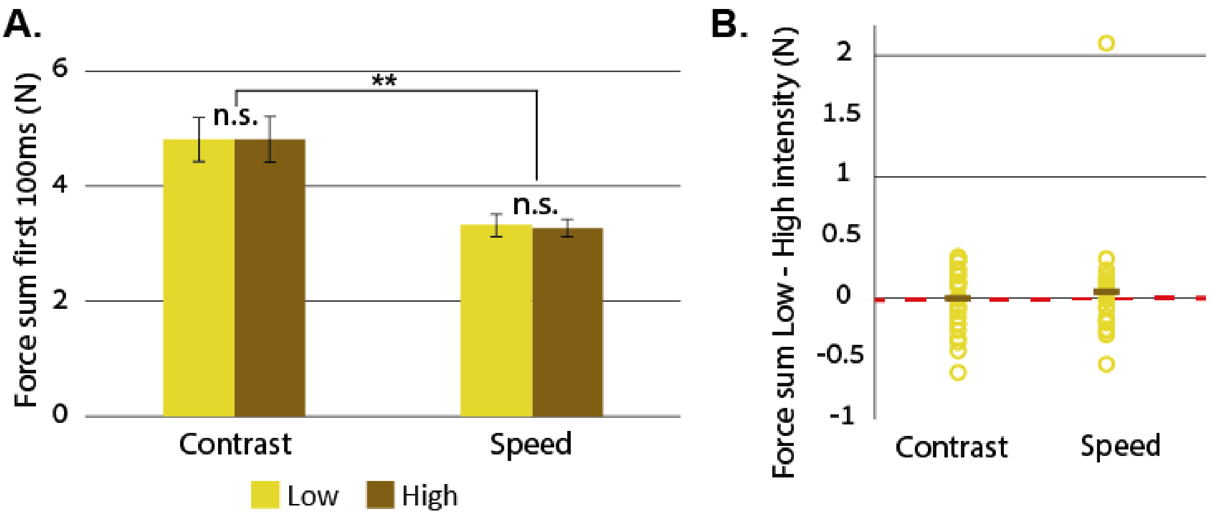


**Figure S1: A.** group average force data at first 100ms of press. ** notes p<0.001. **B.** Individual subjects' differences between low and high intensity outcomes. The dark line represents group average, red line represents equal force for both intensities.

**Summary:**

Taken together, these results further support our previous finding in the visual modality of no difference in press force between contrast levels, even when increasing the perceptual differences between stimuli. Furthermore, these results also demonstrate a lack of effect of movement speed on applied force, even though this could be considered a more 'natural' outcome of visuo-motor coupling. This further implies a functional difference between visuo-motor integrations vs. audio-motor and somato-motor integrations (see discussion).
